# Archetypal Analysis for Population Genetics

**DOI:** 10.1101/2021.11.28.470296

**Authors:** Julia Gimbernat-Mayol, Daniel Mas Montserrat, Carlos D. Bustamante, Alexander G. Ioannidis

## Abstract

The estimation of genetic clusters using genomic data has application from genome-wide association studies (GWAS) to demographic history to polygenic risk scores (PRS) and is expected to play an important role in the analyses of increasingly diverse, large-scale cohorts. However, existing methods are computationally-intensive, prohibitively so in the case of nationwide biobanks. Here we explore Archetypal Analysis as an efficient, unsupervised approach for identifying genetic clusters and for associating individuals with them. Such unsupervised approaches help avoid conflating socially constructed ethnic labels with genetic clusters by eliminating the need for exogenous training labels. We show that Archetypal Analysis yields similar cluster structure to existing unsupervised methods such as ADMIXTURE and provides interpretative advantages. More importantly, we show that since Archetypal Analysis can be used with lower-dimensional representations of genetic data, significant reductions in computational time and memory requirements are possible. When Archetypal Analysis is run in this fashion, it takes several orders of magnitude less compute time than the current standard, ADMIXTURE. Finally, we demonstrate uses ranging across datasets from humans to canids.

**Author summary:** This work introduces a method that combines the singular value decomposition (SVD) with Archetypal Analysis to perform fast and accurate genetic clustering by first reducing the dimensionality of the space of genomic sequences. Each sequence is described as a convex combination (admixture) of archetypes (cluster representatives) in the reduced dimensional space. We compare this interpretable approach to the widely used genetic clustering algorithm, ADMIXTURE, and show that, without significant degradation in performance, Archetypal Analysis outperforms, offering shorter run times and representational advantages. We include theoretical, qualitative, and quantitative comparisons between both methods.

## Introduction

Estimating ancestry cluster allele frequencies and cluster membership from single nucleotide polymorphism (SNP) data is important for many applications in population genetics and applying methods to characterize diverse human cohorts has become an essential part of large-scale genomic studies. With the growing number of samples in whole genome databases, efficient population clustering techniques that can handle such sample sizes have become increasingly important. Existing techniques for the clustering of genomes include STRUCTURE [1], FRAPPE [2] and, ADMIXTURE [3]. These compute probabilistic values referred to as *ancestry coefficients* that represent the fraction of the genome of an individual attributable to a particular population cluster. These methods can perform both supervised and unsupervised inference of *ancestry coefficients*. Supervised inference requires reference individuals from predefined ancestral populations, while unsupervised inference uses the structure of the data alone. These existing approaches perform inference via Bayesian [1] or likelihood based methods [2,3] and tend to be computationally expensive due to the high dimensionality of genomic data.

Dimensionality reduction techniques such as multidimensional scaling (MDS), principal component analysis (PCA) and uniform manifold approximation (UMAP) have been used to overcome the high dimensionality of genomic data [4,5], and have become indispensable for visualization and representation of diversity amongst genomic sequences. In PCA, samples are projected onto the axes of highest variation, each of which is a linear combination of allelic dosages across variants [6]. This method has become particularly important in genome-wide association studies and has also been used to investigate the distribution of genetic variation across geography [7]. An advantage is that no assumptions are made about ancestral populations; however, interpretation can often be misleading if sampling designs are irregular. Unsupervised clustering techniques such as ADMIXTURE or Archetypal Analysis (AA) can complement PCA to provide a detailed description of data and to augment visualization. In this work we show how AA can be coupled with PCA, specifically Single Value Decomposition (SVD), to efficiently cluster samples providing shorter run-times than STRUCTURE or ADMIXTURE. We also discuss how these techniques work, where they differ, and how they relate to well established general-purpose clustering techniques such as K-Means and K-Medioids.

## Materials and methods

### System Overview

The complete proposed pipeline is presented in Figure 1.

**Fig 1.**
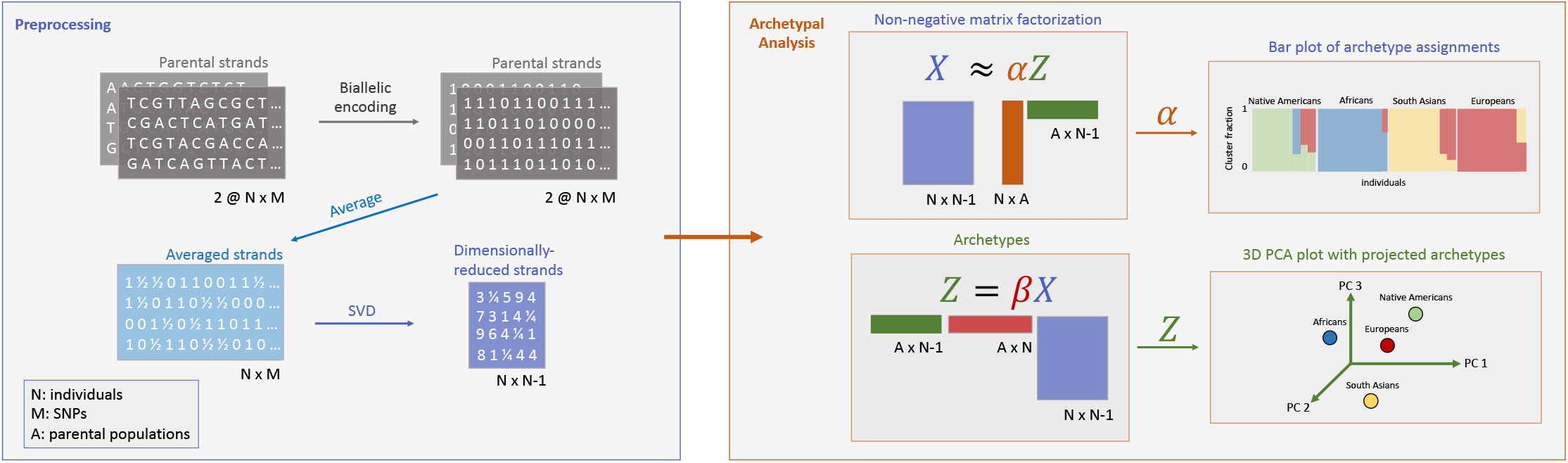
Archetypal Analysis pipeline. The allele counts from both haplotypes of each of *N* individuals are averaged and then dimensionally-reduced from *M* SNPs to *N* – 1 singular vectors via the SVD. Archetypal Analysis then implements an alternating non-negative matrix factorization algorithm that minimizes a constrained sum of squares to find ancestry proportions (*α*) and cluster centroids (*Z*: archetypes). Archetypal analysis models the individual genotypes as originating from the admixture of *A* parental populations, where *A* is an input parameter. For visualization we create bar plots for proportions of archetype assignments given by the matrix *α*, and project archetypes *Z* into a 3D subspace using the first three principal components of the individual genotype sequences.

#### Singular Value Decomposition

Because the subspace spanned by the centered genotype vectors can have no more than *N* – 1 dimensions with *N* the number of samples, there is no loss of information in projecting these centered genotype vectors onto their top N right singular vectors before applying Archetypal Analysis. This operation corresponds simply to a rotation of the coordinate system followed by a pruning of the unused dimensions and yields a space that is generally far smaller than the original, that is the number of total genotyped positions M, since typically *N* ≪ *M*.

If we observe *N* individuals at *M* SNP positions, each individual *i* can be represented by a vector 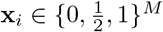, where each position *j* in **x***_i_* indicates the average number of alternate alleles found for each *j* (position) and *i* (individual’s diploid genome). By aggregating **x***_i_* the vectors for all individuals, we obtain an *M* × *N* genotype matrix **G** = [*x*_1_…*x_N_*]. We center the columns of **G** to produce data matrix **X** and then compute the SVD:

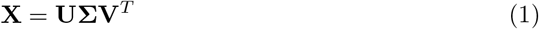

This yields **U** and **V**, the left and right-singular vectors respectively. The first *N* – 1 scores **UΣ** can then be used as input for Archetypal Analysis. As described in [6] these vectors are made up of a linear combination (rotation) of genotypic values across the genome.

#### Archetypal Analysis

This non-negative matrix factorization method was first developed by Cutler and Breiman in 1994 [8], and here it represents each individual as a convex combination of *extreme points*, or archetypes, in allele frequency space. In particular, given an *N* × *M* multivariate data set *X* with *N* individuals and *M* SNPs, for a given number of archetypes or clusters *K*, the algorithm finds the *M* × *K* matrix of archetypes *Z* according to two principles:

1. The samples are approximated as convex combinations of the archetypes such that the the residual sum of squares (RSS) between the approximation and original data is minimized:

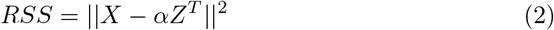

with *α* representing the fractional ancestry assignments, so 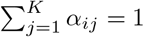, 1 ≥ *α_ij_* ≥ 0 for *i* = 1,…, *N*, and *j* = 1,…, *K*.
2. The archetypes are convex combinations of the samples:

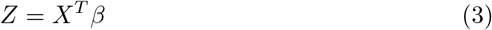

with *β* an *N* × *K* matrix and *β_ij_* indicating the weight of sample *i* at archetype *j*, and 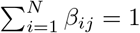 with 1 ≥ *β_ij_* ≥ 0.

By combining Equation 2 and 3 we have:

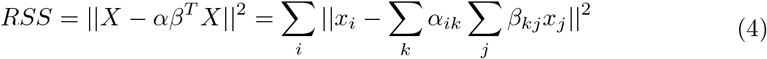

The optimization problem presented in Equation 2 and 3 consists of finding the weight matrices *α* and *β* for a given data matrix *X* and a particular number of archetypes *K*. This is commonly solved through an iterative process of optimizing *α* and *β* in an alternating fashion. For a fixed set of values for *α*, finding the optimal values for *β* is reduced to a constrained least squares problems, and vice versa [8]. The iterative process is typically repeated until the quality of the decomposition reaches a pre-defined threshold, or up to a fixed maximum number of steps. The constrained least square optimization problem can be solved through a variety of techniques. Here we make use of the implementation of [9], which utilizes a non-negative least squares solver obtaining *α_ij_* ≥ 0 and *β_ij_* ≥ 0, where it adds an extra dimension to enforce 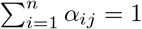 and 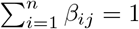. There are multiple open-source packages available in R [10], Python [9] and MATLAB [11] that implement Archetypal Analysis.

Unlike ADMIXTURE, Archetypal Analysis permits the use of dimensionally-rotated representations of SNP data, such as the singular value decomposition. If all singular vectors are used the residual sum of squares of the decomposition (*RSS*’) using projected data *X*’ is equivalent to the *RSS* of the original decomposition:

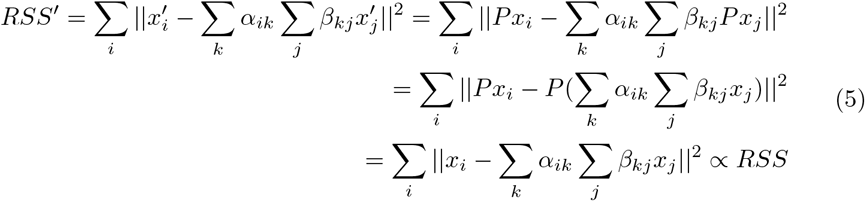

Since the projection matrix *P* = *V*, the orthonormal rotation matrix of *X* onto its singular vector axes.

This permits us to perform AA clustering on a matrix of dimensions only *N* × *N* – 1 instead of *N* × *M*. Note that although the learnt parameters of AA, *α* and *β*, do not depend on *M*, the computation times for *Z* and the *RSS* do, therefore, working in lower dimensions reduces the computational load.

##### Constrained Optimization

Non-negative least squares (NNLS) is a constrained least squares problem in which coefficients are always non-negative (Eq. 8). Archetypal Analysis includes an additional constraint coefficient *C* and adds a row of ones to matrices involved in optimization after every NNLS iteration (Eq. 9 and 10) to ensure the coefficients also sum to one, one of the definitional properties of Archetypal Analysis.

Given an *N* × *M* matrix *X* representing a multivariate data set with *N* observations and *M* attributes, for a given *K*, we minimize:

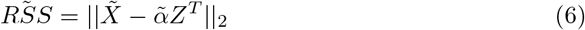

where 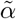 is defined as:

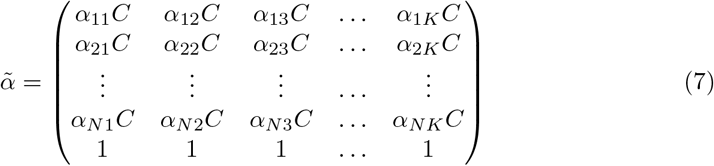

and *α* and archetypes are defined are defined in the previous section. 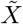 is defined as:

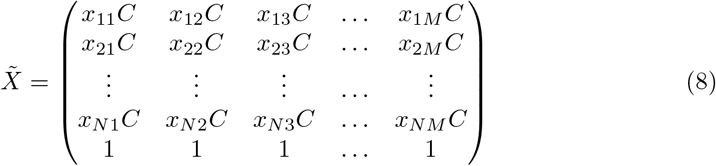

where *C* is a constraint coefficient for *C* > 0 and rows of 1’s are added after every NNLS iteration. This ensures the constraint 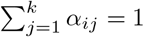 where the value of *C* represents a weighting between the importance of the constraint and NNLS minimization, with lower *C*’s giving a stronger importance to the constraint. The same method is applied to *β* coefficients to ensure 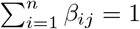.

##### Archetype Initialization

We make use of the implementation in [9] which supports three different archetype initialization strategies: (1) random initialization of the archetypes where each dimension of the archetype is sampled from a uniform distribution scaled to have the same range as the input data, (2) random selection of a sample from the input data as the archetype, and (3) the FurthestSum introduced in [11]. By default we make use of FurthestSum initialization as it efficiently generates initial archetype candidates by, after selecting the first archetype randomly, selecting each subsequent archetype as the sample that has the largest aggregate distance from the previously selected archetypes.

##### Implementation Details

Archetypal analysis was run with the following parameters (with code adapted from [9]).

- Tolerance: defines when to stop optimization when alternating between finding the best *α*’s for given archetypes *Z* and finding the best *Z* for given *α*’s. Specifically, the threshold applied is,

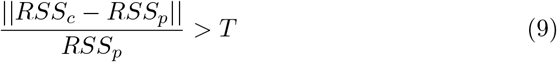

where *RSS* is the residual sum of squares defined in (Eq. 2) for the current iteration *RSS_c_* and the previous iteration *RSS_p_*, and *T* is the desired tolerance. We use a value of *T* = 0.001.
- Maximum number of iterations for the residual sum of squares (*RSS*) minimization: 50.
- Constraint coefficient *C*: coefficient that ensures the summation of *α*’s and *β*’s equals to 1. See Appendix B for further details on the constrained optimization method. We use a value of *C* = 0.001.
- Initialization method: we use FurthestSum [11] as the initialization method.

### Datasets

#### HUMAN

Whole genomes from the Human Genome Diversity Project [12], the Simons Genome Diversity Project [13] and the 1000 Genomes Project [14] have been included in this study. The Human Genome Diversity Project whole genome cohort includes 929 individuals from 54 human populations. The Simons Genome Diversity Project contains 300 genomes from 142 diverse populations, and the 1000 Genomes Project includes 2504 individuals from 26 populations. The three datasets were merged, removing duplicated individuals between the studies and retaining only SNPs present in all three datasets, yielding an intersection of 1,411, 471 SNPs for analysis. Rare variants with minor allele frequencies < 0.1 were removed. In total, 3558 individuals were included in the study from 7 different continents: 683 from Europe, 805 from Africa, 34 from Oceania, 695 from South Asia, 772 from East Asia, 150 from West Asia, and 419 indigenous individuals from the Americas.

#### Dogs

The heterogeneous data set of dog breeds from [15] consists of 1355 groups representing 166 dog breeds. Each sequence has a total of 150, 131 SNPs. Populations with vastly different histories are included, originating from all continents except Antarctica [15].

## Results

### Human datasets

#### Principal Components and Archetypal Analysis

We first compute the principal components of the human data set and display the first two components in a plot coloured by continental population (Fig. 2, **a**). The African population displays the highest genetic variability extending across the first principal component axis (11% explained variance). We then use all principal components, that is the projection onto all the left singular vectors of the SVD, as input to the Archetypal Analysis and plot the proportional membership of each cluster for each individual in a compositional plot (Fig. 2, **b**). The African population is represented by three archetypes (A1, A2 and A8), while the East Asian and South Asian populations have one archetype each (A3 and A5 respectively). Note that Archetypal Analysis addresses the high variation within African groups by using multiple archetypes. The European and West Asian populations share a single archetype (A4), while the Oceanian populations are found on the gradient between the East Asian and South Asian archetypes. Finally, the Native American populations are represented by two archetypes (A6 and A7) and have a gradient running to the European/West Asian archetype due to colonial admixture. Example populations found along this gradient are the Puerto Ricans and Colombians.

**Fig 2.**
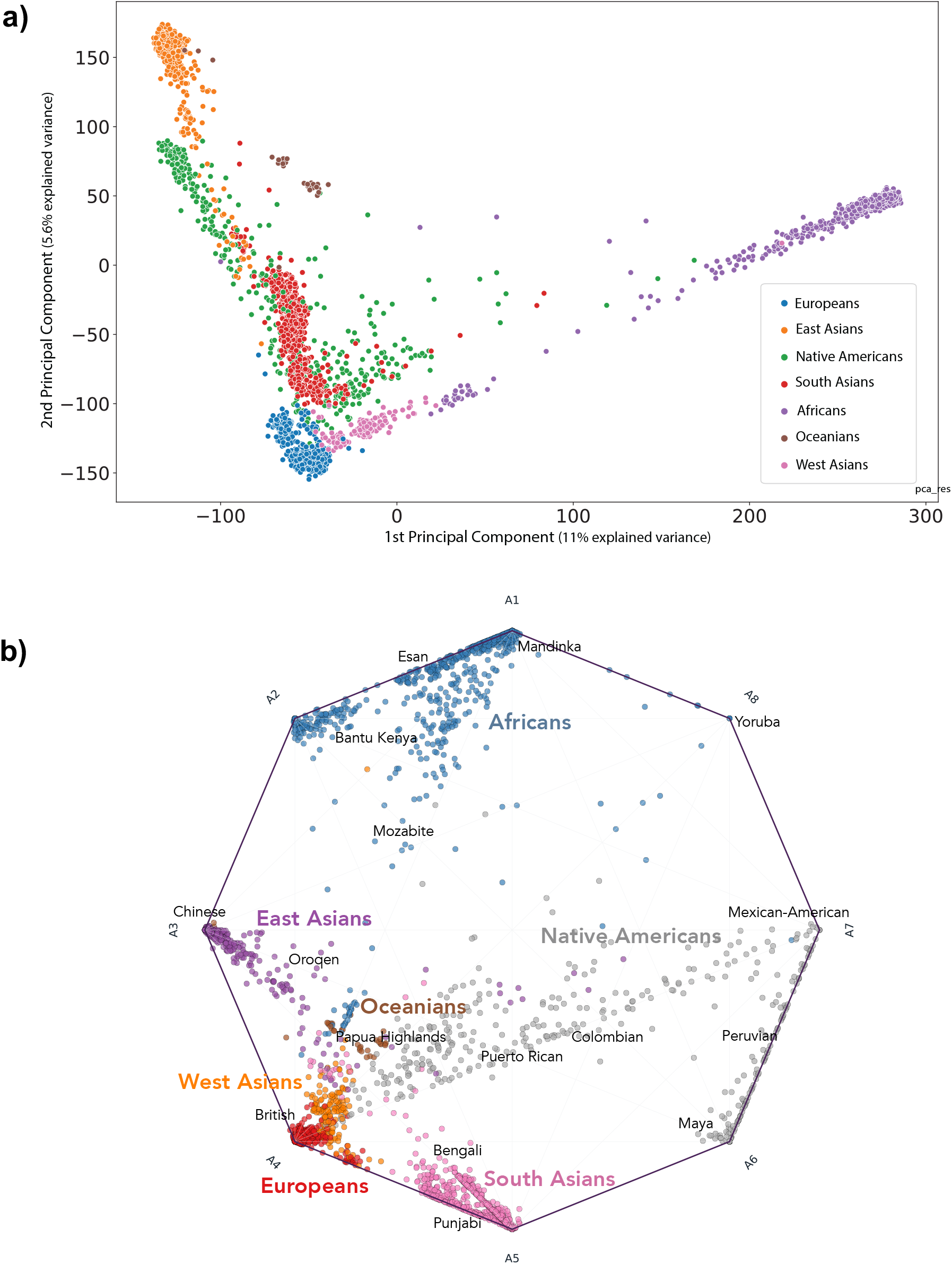
Principal Component Analysis and Archetypal Analysis compositional plots for human populations (K=8). **a)**, 2-dimensional PCA plot of human continental populations, where groups of individuals are colored by the unique regional genetic components they possess (see legend) **b)**, Compositional plot giving proportional archetype assignment for each individual (points). Points are coloured by the presence of regional genetic components and a few example sub-populations are shown given in text. Clusters of individuals from the same population are observed on the edges of the polygon and gradients between edges indicate admixed individuals.

#### Comparison of ancestry estimates

To compare the ancestry estimates derived from ADMIXTURE and Archetypal Analysis, we display the proportional ancestry cluster assignments, the *Q* and *α* matrices respectively, in a bar plot for *K* = 8 cluster (Fig. 3, **b**). Each vertical bar represents an individual and the shaded colors denote the cluster proportions. We also display individuals on a three-dimensional PCA plot with projected archetypes (*Z*) and ADMIXTURE cluster centers (*F*) (Fig. 3, **a**). A theoretical comparison of both methods can be found in the Discussion section.

**Fig 3.**
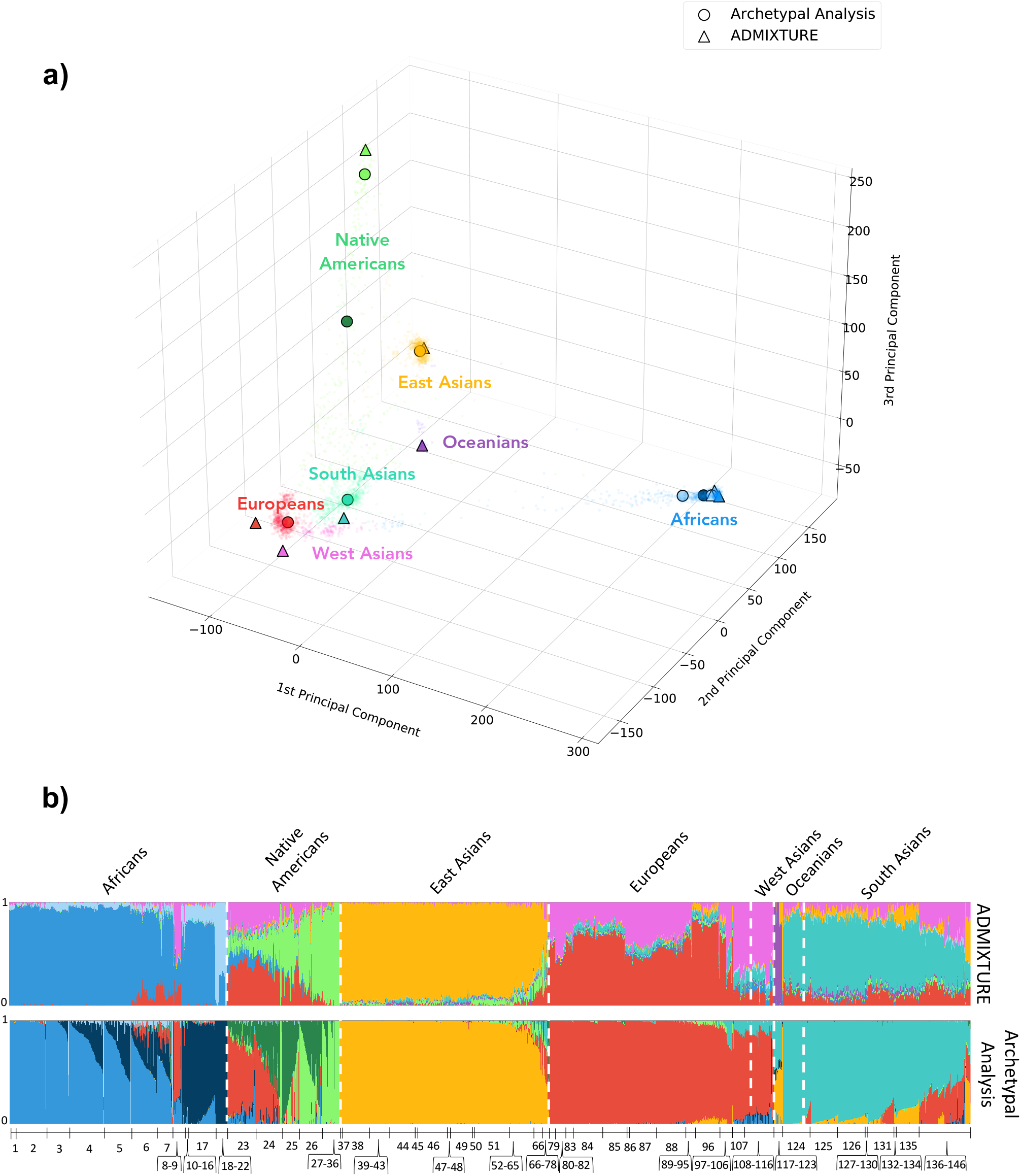
Comparison of ancestry estimates for human populations (K=8). **a)**, three-dimensional PCA plot of individuals with projected archetypes (circles) and ADMIXTURE cluster centers (triangles). **b)**, bar plot where individual are represented along the horizontal axis as narrow columns and ordered by population group. Colour bars along the vertical axis show the proportional cluster assignment for each individual. We compare the cluster assignments of ADMIXTURE (top) and Archetypal Analysis (bottom).

##### Archetypal Analysis

European (red), South Asian (turquoise), and East Asian populations (yellow) are predominantly represented by a single archetype. Native American populations are a combination of three archetypes, two of which are mostly specific to this population (light green and dark green) and a third, representing colonial admixture, which is European (red). Individuals from Puerto Rico and Colombia mostly share the third archetype with Europeans. The African population is represented by three archetypes. One archetype encompasses West African populations such as Mandeka, Gambian Mandika and Mende (ocean blue). Another includes eastern and southern groups such as Luhya and San (navy blue). A third archetype represents a few individuals from all African populations (light blue).

##### ADMIXTURE

Oceanian (purple) and East Asian populations (yellow) are predominantly represented by a single cluster center. Europeans and West Asians are a combination of two centers (red and pink) that are located outside the point cloud of individuals, differing from AA which captures both with a unique cluster. Native Americans show traces of the European and West Asian cluster components, but are mostly represented by their own, here single, cluster (light green). African populations are predominantly represented by two clusters (ocean blue and light blue), while a few populations, such as the North African Mozabites, show traces of European and West Asian components. Finally, South Asians predominantly cluster around a single cluster (turquoise), but also show traces of the European and West Asian clusters.

Overall, Archetypal Analysis provide estimates which qualitatively often match ethnolinguistic and geographical labels. AA properly captures the wide variation within African populations; however, it fails to identify a unique cluster for Oceanians. Additionally, due to its stronger constraints than ADMIXTURE, AA obtains cluster centroids that lie near actual sampled genotypes.

### Domestic dog breed dataset

#### Principal Components and Archetypal Analysis

We compute the principal components of the dog breed data sets and display the first two components in a plot coloured by dog clades (Fig. 4, **a**). The Asian Spitz clade shows the highest genetic variability extending across the first principal component axis, including breeds such as Chow Chow, Greenland Sledge Dog and Siberian Husky. The latter is found close to the wolf, while the European Mastiff clade represented by breeds such as Bull Terrier, Boxer and Bulldog extends across the second principal component axis. Archetypal Analysis is then computed for *K* = 5 and *K* =15 with principal components as input (Fig 4, **b** and **c**). For *K* = 5, dog archetypes were found to be the Asian Spitz dogs (A1), the Bulldog-derived dogs (A2), the Terriers (A3), hunting water dogs (A4) and herding dogs (A5). The remaining breeds are displayed as a combination of these main archetypes, mostly represented by A5 and A4. This matches the structure shown in PCA, where most of the breeds are clustered in the origin, except the dogs in the Bull Terrier and Husky groups. When increasing the number of archetypes to *K* = 15, individual dog breeds begin clustering around single archetypes, showing the growing population structure. New archetypes appear for the Boxer (A3), Irish Wolfhound (A4), Otter Hound (A5), Bullmastiff (A6), Bernese Mountain Dog (A10), Glen of Imaal Terrier (A11), French Bulldog (A12), Boston Terrier (A13), Shetland Sheepdog (A14) and Tibetan Spaniel (A15). The rest of the breeds are mostly found near A14 and A15.

**Fig 4.**
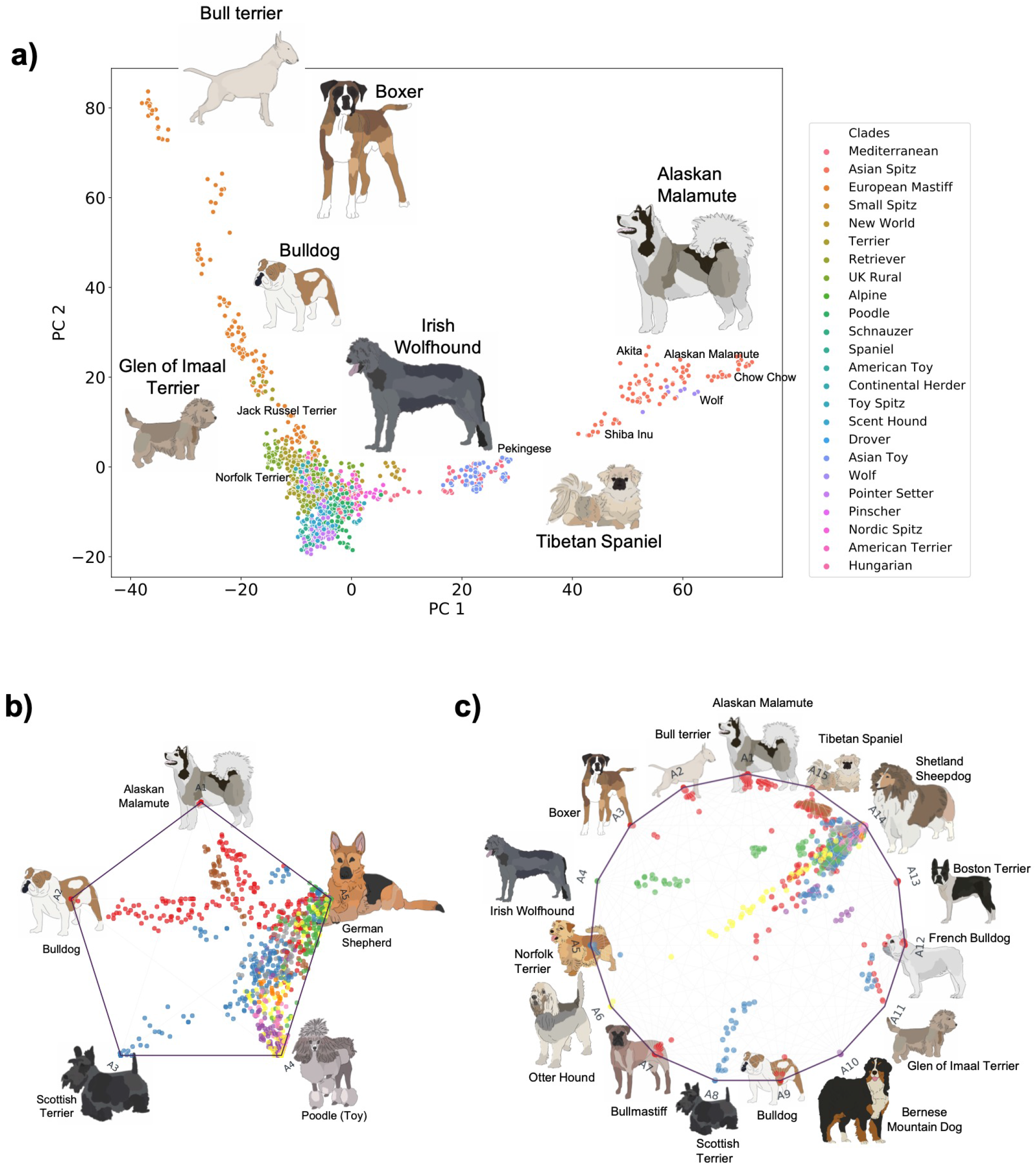
Principal Component Analysis and Archetypal Analysis compositional plots for domestic dog breeds. **a)**, two-dimensional PCA plot of domestic dog breeds where groups of dogs are colored by clade. **b)** and **c)**, proportional composition of each cluster for each individual in coordinate space for K=5 and K=15 respectively. Data points are coloured by clade and archetype representatives are shown as drawings. Gradients between edges indicate combinations between breeds.

#### Performance metric analysis

The dog breed dataset was used to benchmark the computation times and clustering quality of both ADMIXTURE and Archetypal Analysis. Running times and explained variances of ADMIXTURE and Archetypal Analysis are measured for an increasing number of archetypes/clusters *K* = 1,…, 22 and *K* = 1,…, 30 respectively. The initialization was set to *random* for both methods to achieve uniform comparison and results were averaged over 5 runs. Accumulated run-times increased exponentially with *K* for ADMIXTURE whereas they increased linearly for Archetypal Analysis (Fig. 5). An accumulated runtime of 34 minutes was taken by Archetypal Analysis to compute ancestry estimates for *K* = 2 to *K* = 30 clusters. For ADMIXTURE, the accumulated runtime from *K* = 2 to *K* = 30 was 78 hours. Thus, Archetypal Analysis ran 137 times faster than ADMIXTURE on the domestic dog breed dataset. A similar increase in relative speed was maintained, on average, for non-cumulative times (Table 1).

**Fig 5.**
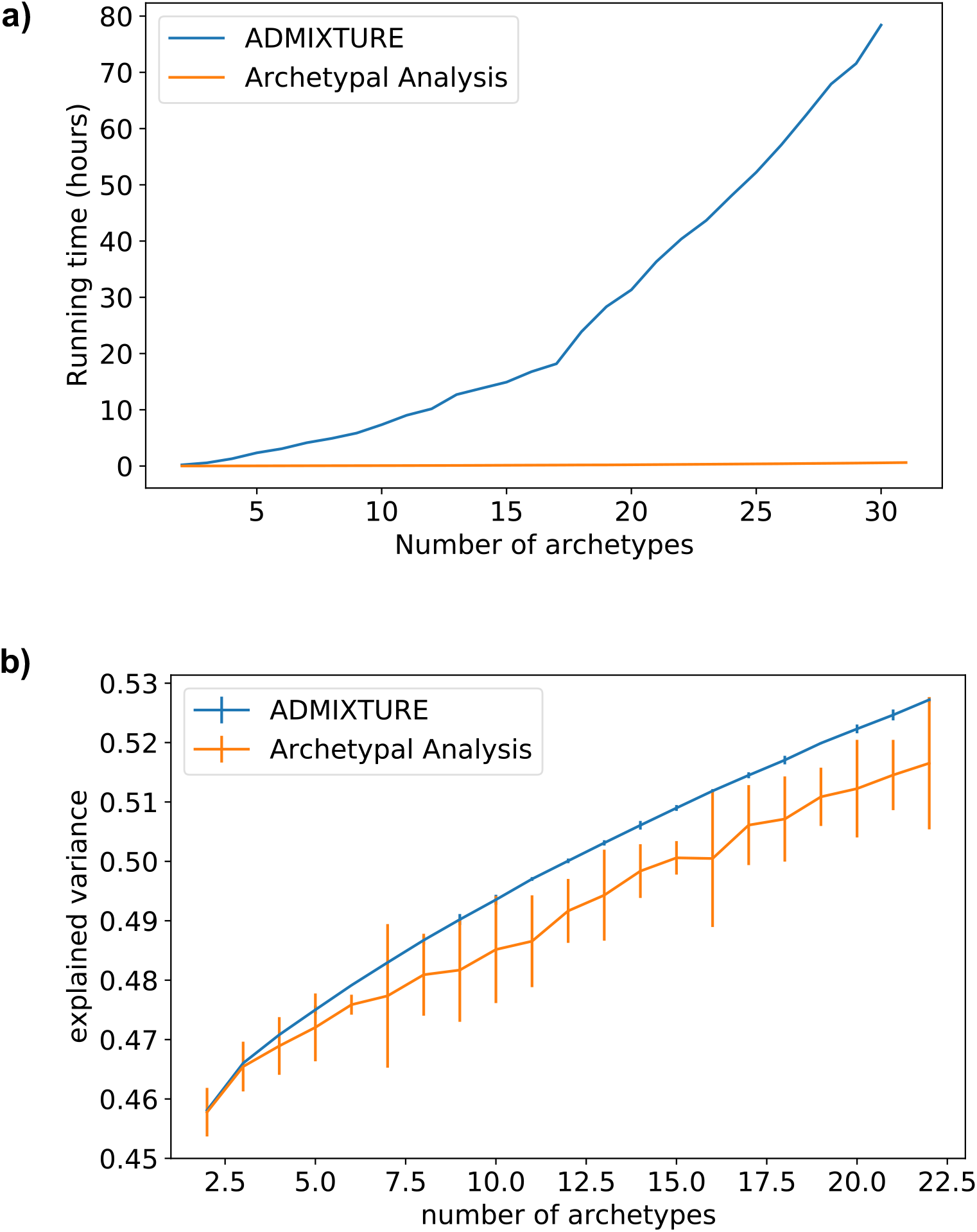
Performance metrics analysis. **a)**, runtime analysis for ADMIXTURE and Archetypal Analysis for *K* = 2 to *K* = 30. Time is expressed in units of accumulated hours. **b)**, explained variance analysis comparison for ADMIXTURE and Archetypal analysis for *K* = 2 to *K* = 22. Results are averaged over five distinct random seed values for each value of K and the ranges observed are shown as vertical bars.

**Table 1.** Runtime (in minutes) for ADMIXTURE-AA comparison.

|  | K (number of clusters / archetypes) |  |  |  |  |  |  |
| --- | --- | --- | --- | --- | --- | --- | --- |
| Algorithm | [2-6) | [6-10) | [10-14) | [14-18) | [18-22) | [22-26) | [26-30) |
| ADMIXTURE | 43 | 64 | 97 | 150 | 247 | 250 | 319 |
| AA | 0.5 | 0.48 | 0.7 | 1 | 1.4 | 1.9 | 2.4 |
| Relative speed | 86× | 133× | 139× | 150× | 176× | 132× | 132× |

**Table 2.** ADMIXTURE and Archetypal Analysis comparison.

|  | ADMIXTURE | Archetypal Analysis |
| --- | --- | --- |
| **Model** | $X \approx QF$ | $X \approx \alpha Z^T$ |
| Loss Function | log-likelihood | RSS |
| Free-parameters | $(N + M)K - N$ | $2NK - N - K$ |
| **Cluster Assignments (CA)** | $Q$ | $\alpha$ |
| CA Dimensions | $N \times K$ | $N \times K$ |
| CA Free-parameters | $N(K - 1)$ | $N(K - 1)$ |
| CA Constraints | $\sum_{j=1}^K Q_{ij} = 1$ and $Q_{ij} \geq 0$ | $\sum_{j=1}^K \alpha_{ij} = 1$ and $\alpha_{ij} \geq 0$ |
| **Cluster Centroids (CC)** | $F$ | $Z = X^T \beta$ |
| CC Dimensions | $K \times M$ | $K \times M$ |
| CC Free-parameters | $KM$ | $K(N - 1)$ |
| CC Constraints | $0 \leq F_{ij} \leq 1$ | $\sum_{j=1}^N \beta_{ij} = 1$ and $\beta_{ij} \geq 0$ |

Explained variances increased linearly in the number of clusters for both algorithms (Fig. 5). The explained variance for Archetypal Analysis was on average 2% lower than for ADMIXTURE. For the values of *K* included in this analysis, the mean standard deviation for five averaged runs with random initialization was 0.007 for Archetypal Analysis and 0.0004 for ADMIXTURE. As described in the following Discussion section, the difference in explained variance is due, at least in part, to the stronger restrictions that Archetypal Analysis imposes when estimating the cluster centroids. However, as shown with human sequences in figure 3, the stronger restrictions of AA lead to centroids that are always a linear combination of actual samples, guaranteeing that they represent theoretically observable population samples.

## Discussion

### Population structure overview

Archetypal Analysis proved to be an interpretable alternative to ADMIXTURE. It assigned separate regional archetypes that associated predominantly with Europeans, with South Asians, and with East Asians, and it recognized the high genetic variability of African populations. Differences within regions were also detectable (Fig. 3). For example, indigenous peoples across the Americas were separated from the remainder of the modern American communities as the light green archetype. Peruvians were also included in this group, most likely because indigenous groups make up 45% of the Peruvian population. Similarities in peoples that are geographically spread were also detected. For example, the Bantu peoples (Bantu Herero, Bantu Tswana, Bantu Kenya, Bantu South Africa and Luhya) comprise several hundred indigenous ethnic groups in Africa spread over a vast area from Central Africa to Southern Africa, but those present in our dataset were grouped together forming the dark blue archetype.

European-like archetype components seen in African peoples due to geographic proximity and migration were also found in the Saharawi and Mozabites from the northwestern part of Africa. As also observed in previous studies, American populations, such as Puerto Rico and Colombia, showed European representation due to Spanish colonization. The suggested effects of this historical event can also be observed in (Fig. 2, b), which shows a gradient of relatedness to Europeans that runs through Puerto Ricans, Colombians, Peruvians through the Mexican-Americans. Archetypal Analysis also identified South Asian communities having a shared component with Europeans that ADMIXTURE did not detect (Fig. 3, b). For example, the Brahui, Kalash and Baloch were identified with a European-like archetype by Archetypal Analysis and not by ADMIXTURE. These might reflect the influence of Indo-European migrations and the Ancestral North Indians [16], an ancestral genetic grouping in India that shares some ancestry with other Indo-European speakers from India to Iran to Europe.

### Relationship between Archetypal Analysis and ADMIXTURE

The popular algorithm ADMIXTURE estimates individual ancestries by computing maximum likelihood estimates in a parametric model. Specifically, it maximizes the biconcave log-likelihood of the model using block relaxation:

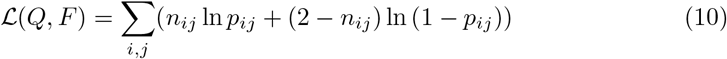

where genotype *n_ij_* for individual *i* at SNP *j* represents the number of type ‘1’ alleles observed. Given *K* populations, the success probability 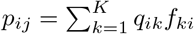 in the binomial distribution *n_ij_* ~ Bin(2,*p_ij_*) depends on the fraction *q_ik_* of *i*’s ancestry attributable to population *k* and on the frequency *f_kj_* of the allele 1 in population *k*, where *q_ik_* and *f_kj_* are the entries of *Q* and *F* respectively [3].

ADMIXTURE and Archetypal Analysis share similar modeling assumptions. Both *q_kj_* ADMIXTURE and *α* archetype fractions can be interpreted as partial cluster assignments while ADMIXTURE frequency coefficients *f_kj_* and archetype coordinates *Z* encode cluster center locations in SNP space. A key difference is that ADMIXTURE cluster centroids have M (# of SNPs) free parameters, in other words, the frequency at each SNP for each cluster (*f_kj_*) is a parameter that needs to be learnt. Instead, in AA, cluster centroids have N (number of samples) free parameters, that is, a coefficient (*β*) for each training sample needs to be learnt for each cluster center. When M ≫ N (the typical scenario when working with genomic data), AA has far fewer free-parameters than ADMIXTURE. This can lead to lower explained variance values (or higher reconstruction errors), but guarantees centers that exist within the convex hull of real samples (and thus could represent a real descendant individual), while ADMIXTURE can over-fit, yielding centers outside the hull of the observed data (see Results section) that may represent no population that has ever existed. Furthermore, because AA does not optimize each of the M free-parameters, it can work with rotated data (the left singular vectors of the SVD) without any loss of information, or with dimensionally-reduced data, allowing for a much more efficient computation.

The likelihood function of ADMIXTURE can be understood as an error or distance metric between the input sequences *X* (where both haplotypes have been averaged) and a decomposed product *QF*. In fact, when *X* ≈ *QF*:

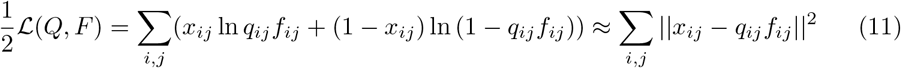

Therefore, the likelihood function resembles the *RSS* problem of AA. In fact, ADMIXTURE can be understood as a type of likelihood-based relaxed archetypal analysis, where the constraints imposed on the cluster centroids are loosened.

Another shared aspect of both methods is the alternating nature of the optimization procedure. In both methods, cluster centers and cluster assignments are optimized in an iterative manner. Once the cluster assignments are fixed, optimizing centers becomes a convex problem, and vice versa, allowing for fast convergences.

### Relationship between Archetypal Analysis, ADMIXTURE, K-Means, and K-Medioid Clustering

Archetypal Analysis and ADMIXTURE hold a strong relationship with K-Means and K-Medioids. As already stated in [11], if the constraints on the archetypes *Z* are relaxed, and cluster assignments are limited to binary values *α* ∈ {0,1} and 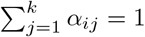, then archetypal analysis becomes equivalent to K-Means. Similarly, if the sparsity regularization used in ADMIXTURE [3] is strongly applied, the clusters assignments Q become binary and the technique becomes similar to K-Means. In a similar fashion, if both *α* and *β* are restricted to be binary, *α*, *β* ∈ {0, 1}, Archetypal Analysis becomes equivalent to K-Medioids. Therefore, AA can be understood as a smooth or fuzzy version of K-Medioids. Note that both K-Means and K-Medioids are also typically optimized in a iterative alternating nature, similar to AA and ADMIXTURE.

Figure 6 shows a qualitative comparison of all four of these methods when *K* = 4. Examples with *K* = 3 and *K* = 5 can be found in the supplement. We can observe that ADMIXTURE with sparsity constraints (green) obtains cluster centroids less extremal than ADMIXTURE without these constraints, showing a behaviour that tends to K-Means. Note that the differences between cluster centers will not depend only on differences in modelling assumptions for each technique, but also in differences in implementation details and initialization approaches of each method.

**Fig 6.**
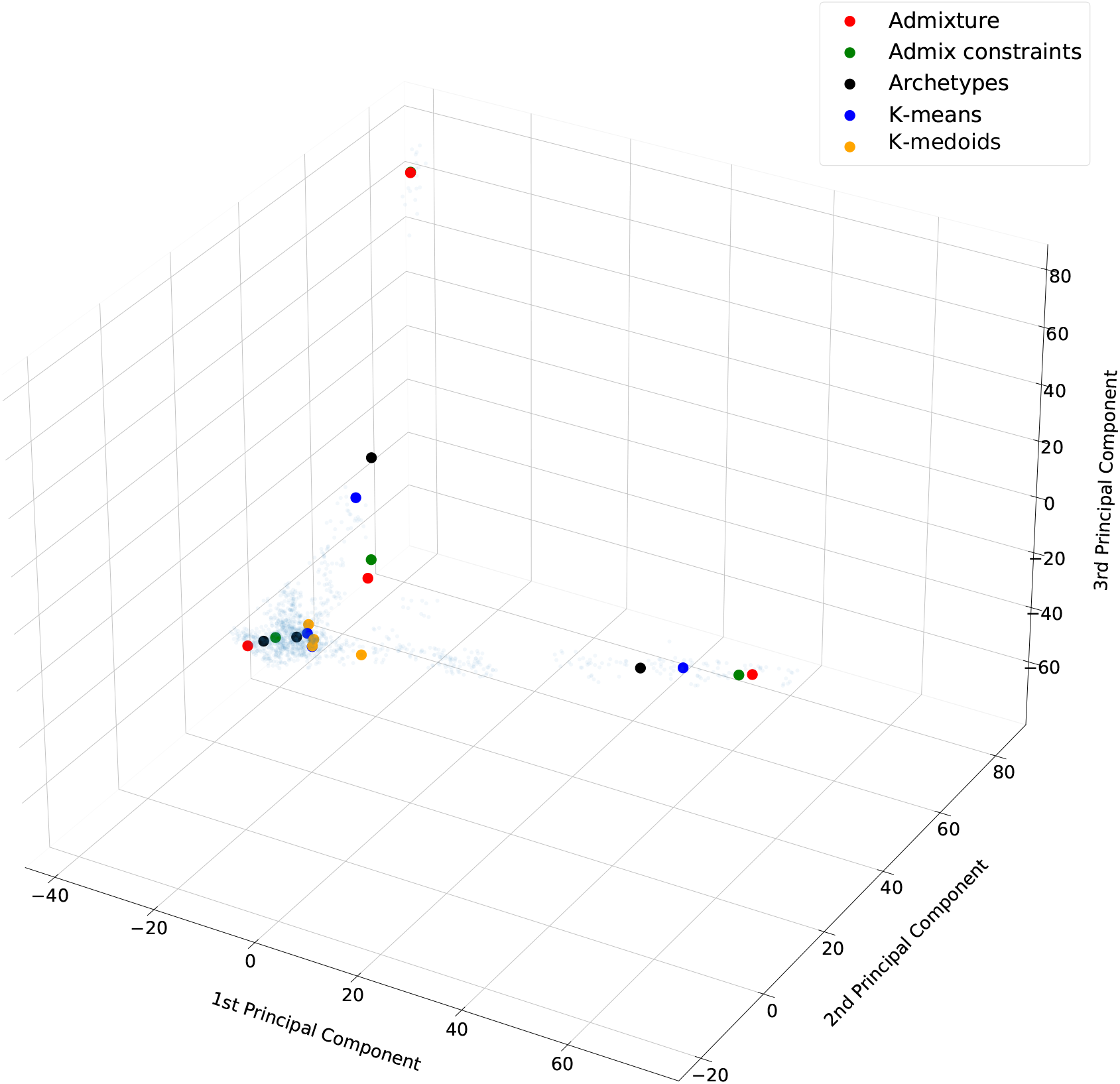
Comparison of cluster centroids from different methods. Cluster centers learned by ADMIXTURE, ADMIXTURE with sparsity regularization, Archetypal Analysis, K-Means, and K-Medoids

## Conclusion

In this paper we show how Archetypal Analysis can be used as a fast alternative to

ADMIXTURE for population clustering. We also show how the Archetypal Analysis model has fewer degrees of freedom, restricting the centroids of clusters within convex hull combinations of the training samples, which leads to lower explained variance than ADMIXTURE, but provide more interpretable cluster centroids. We apply our proposed system to human and dog genotypes, showing that AA can perform more than two orders of magnitude faster than ADMIXTURE while still properly capturing the population structure of the data.

## Acknowledgments

This work was supported in part by the Chan Zuckerberg Biohub (through CDB) and by the Royal Academy of Engineering Leaders Scholarship (awarded to JGM). We would like to thank Santiago De Vilallonga for his illustrations of dogs.

## Supporting information

### Human bar plot labels

Table 3 displays all the subpopulation labels used in Figure **3b**. The details of the dataset can be found in the Datasets subsection in the Methods section.

**Table 3.**
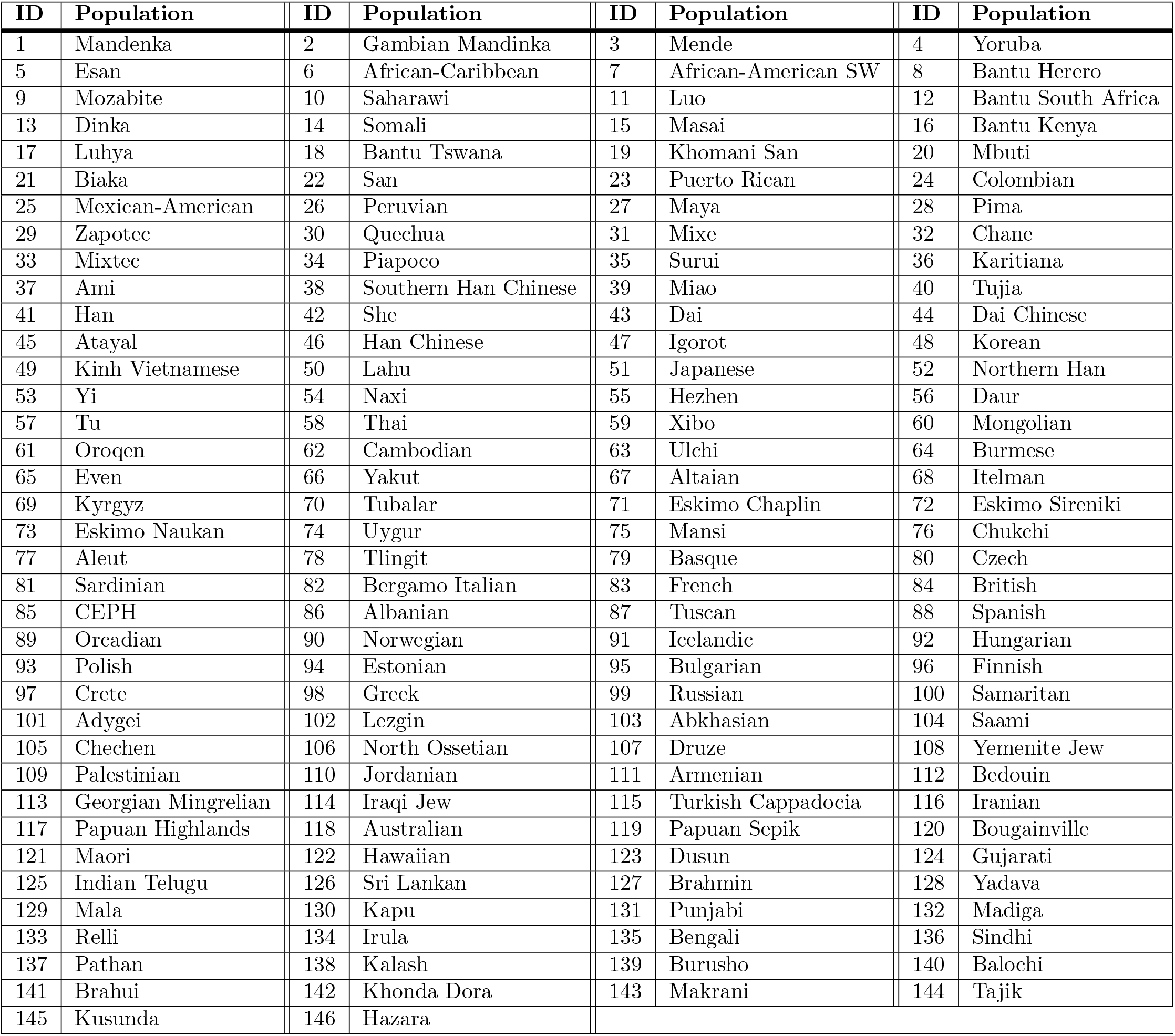
Bar plot labels.

### Domestic dog breeds details

Tables 4 and 5 show the breeds of all the dogs included in our study. Details about the dataset can be found above in the Datasets subsection within the Methods section.

**Table 4.**
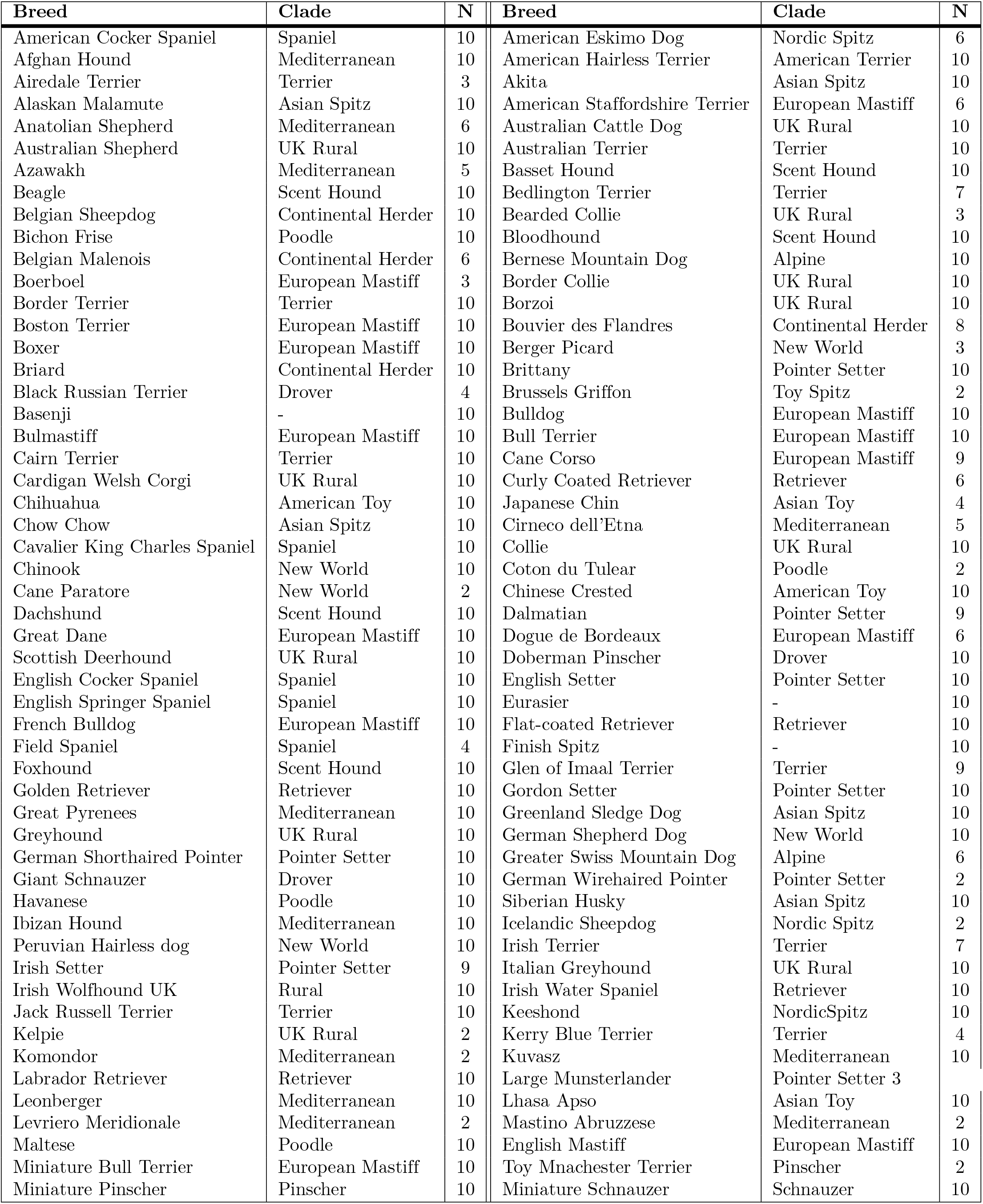
Domestic dog breeds details (1)

**Table 5.** Domestic dog breeds details (2)

| Breed | Clade | N | Breed | Clade | N |
| --- | --- | --- | --- | --- | --- |
| Neapolitan Mastiff | European Mastiff | 6 | Chinese Shar-pei | Asian Spitz | 10 |
| Norwegian Elkhound | Nordic Spitz | 10 | Shiba Inu | Asian Spitz | 8 |
| Newfoundland | Retriever | 10 | Shih Tzu | Asian Toy | 10 |
| Norfolk Terrier | Terrier | 10 | Silky Terrier | Terrier | 4 |
| Norwich Terrier | Terrier | 10 | Schipperke | Toy Spitz | 10 |
| Nova Scotia Duck Tolling Retriever | Retriever | 10 | Sloughi | Mediterranean | 5 |
| Old English Sheepdog | UK Rural | 10 | Spinone Italiano | Pointer Setter | 2 |
| Otter Hound | Scent Hound | 9 | Shetland Sheepdog | UK Rural | 10 |
| Papillon | Toy Spitz | 10 | Standard Schnauzer | Schnauzer | 10 |
| Parsons Russell Terrier | Terrier | 2 | Staffordshire Bull Terrier | European Mastiff | 10 |
| Petit Basset Griffon Vendéen | Scent Hound | 10 | Saint Bernard | Alpine | 10 |
| Pekingese | Asian Toy | 10 | Swedish Valhund | Nordic Spitz | 6 |
| Pembroke Welsh Corgi | UK Rural | 10 | Tibetan Mastiff | Asian Spitz | 10 |
| Pharaoh Hound | Mediterranean | 2 | Tibetan Spaniel | Asian Toy | 10 |
| Pomeranian | Small Spitz | 10 | Tibetan Terrier | - | 10 |
| Poodle - Miniature | Poodle | 10 | Belgian Tervuren | Continental Herder | 10 |
| Poodle - Standard | Poodle | 10 | Toy Fox Terrier | American Terrier | 4 |
| Poodle - Toy | Poodle | 10 | Vizsla | Pointer Setter | 7 |
| Portuguese Water Dog | Poodle | 10 | Volpino Italiano | Small Spitz | 4 |
| Pug Dog | Toy Spitz | 10 | Weimaraner | Pointer Setter | 10 |
| Puli | Hungarian | 4 | Wire Fox Terrier | Terrier | 10 |
| Pumi | Hungarian | 5 | Whippet | UK Rural | 10 |
| Rat Terrier | American Terrier | 2 | Wirehaired Pointing Griffon | Pointer Setter | 6 |
| Redbone Coonhound | Scent Hound | 2 | West Highland White Terrier | Terrier | 10 |
| Rhodesian Ridgeback | European Mastiff | 9 | Xigou | Asian Spitz | 5 |
| Rottweiler | Drover | 10 | Xoloitzcuintle | New World | 5 |
| Saluki | Mediterranean | 19 | Xoloitzcuintle - Miniature | New World | 5 |
| Samoyed | - | 10 | Yorkshire Terrier | Terrier | 10 |
| Scottish Terrier | Terrier | 10 | Grey Wolf | - | 7 |
| Soft Coated Wheaten Terrier | Terrier | 4 | Golden Jackal | - | 2 |

### Archetypal Analysis compositional plots

Figures 7 and 8 show further examples of Archetypal Analysis predictions with the dataset of dog genotypes. The figures show that as the number of archetypes increases, more breeds are clustered in their individual archetype (e.g. A6 to A15 in the 15 archetypes plot), while the rest of breeds (the majority of the breeds) are represented as a combination of a few number of archetypes (e.g. A1 to A5 in the 15 archetypes plot).

**Fig 7.**
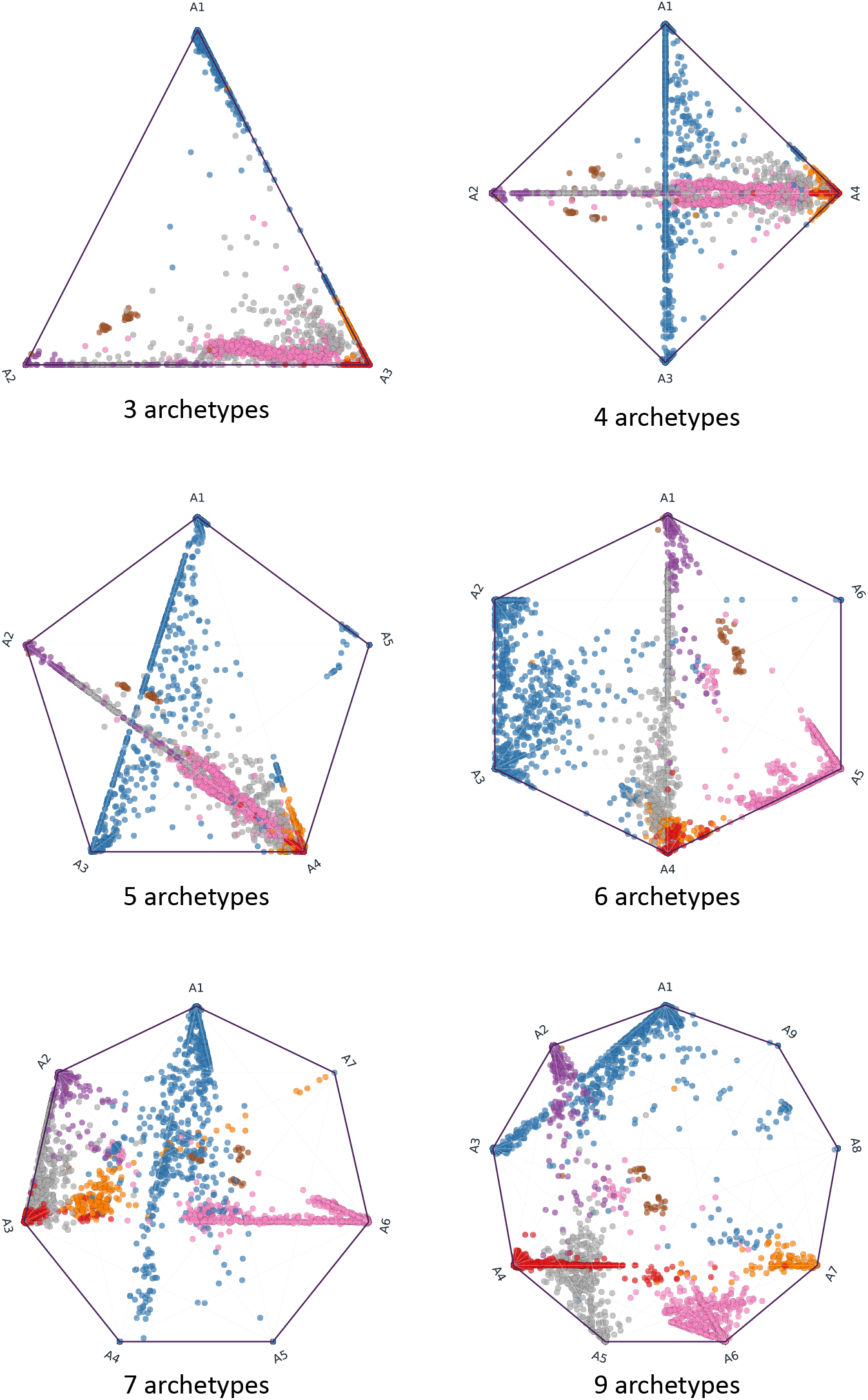
Archetypal Analysis compositional plots for human continental populations. Archetypal Analysis polygon compositions of human data (3-9 archetypes, excepting the 8-archetype polygon which can be found as Fig. 2 in Section 4.1.1). The colours represent the continental origins: EUR - European (red), AFR - African (blue), EAS - East Asian (purple), WAS - West Asian (orange), OCE - Oceanian (brown), SAS - South Asian (pink), AMR - American (gray).

**Fig 8.**
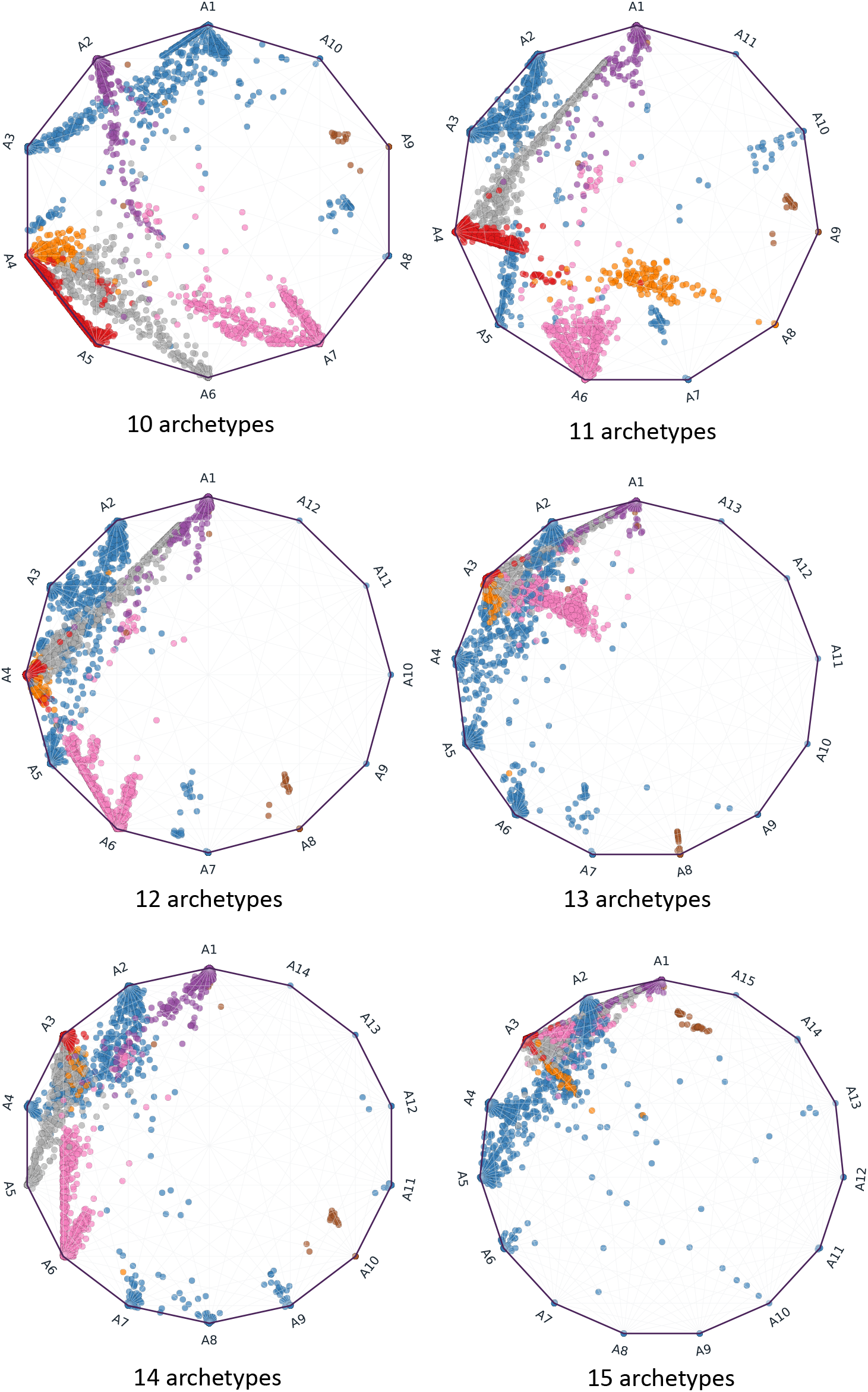
Archetypal Analysis compositional plots for human continental populations. Archetypal Analysis polygon compositions of human data (10-15 archetypes). The colours represent the continental origins: EUR - European (red), AFR - African (blue), EAS - East Asian (purple), WAS - West Asian (orange), OCE - Oceanian (brown), SAS - South Asian (pink), AMR - American (gray).

### Genome-Wide Association Studies

In order to depict how Archetypal Analysis can be included in GWAS, we display a Manhattan plot (Figure 9) and a Q-Q plot (Figure 10) of an association study of the height of dog breeds with and without archetype proportions as covariates.

**Fig 9.**
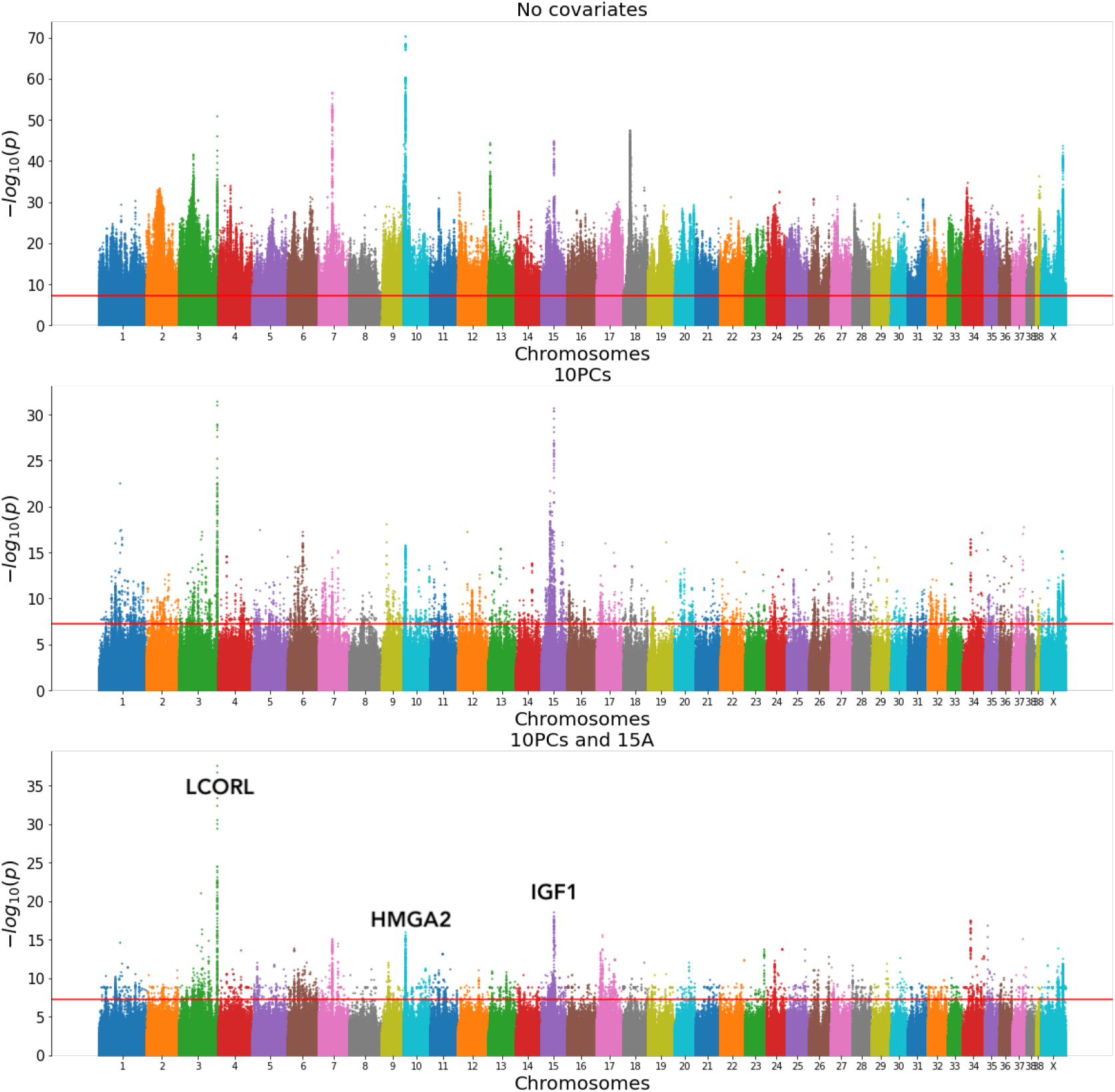
Manhattan plot. Manhattan plots for no covariates (top), when adding 10 PCs (middle) and when adding 10 PCs and 15 Archetype coefficients (bottom).

**Fig 10.**
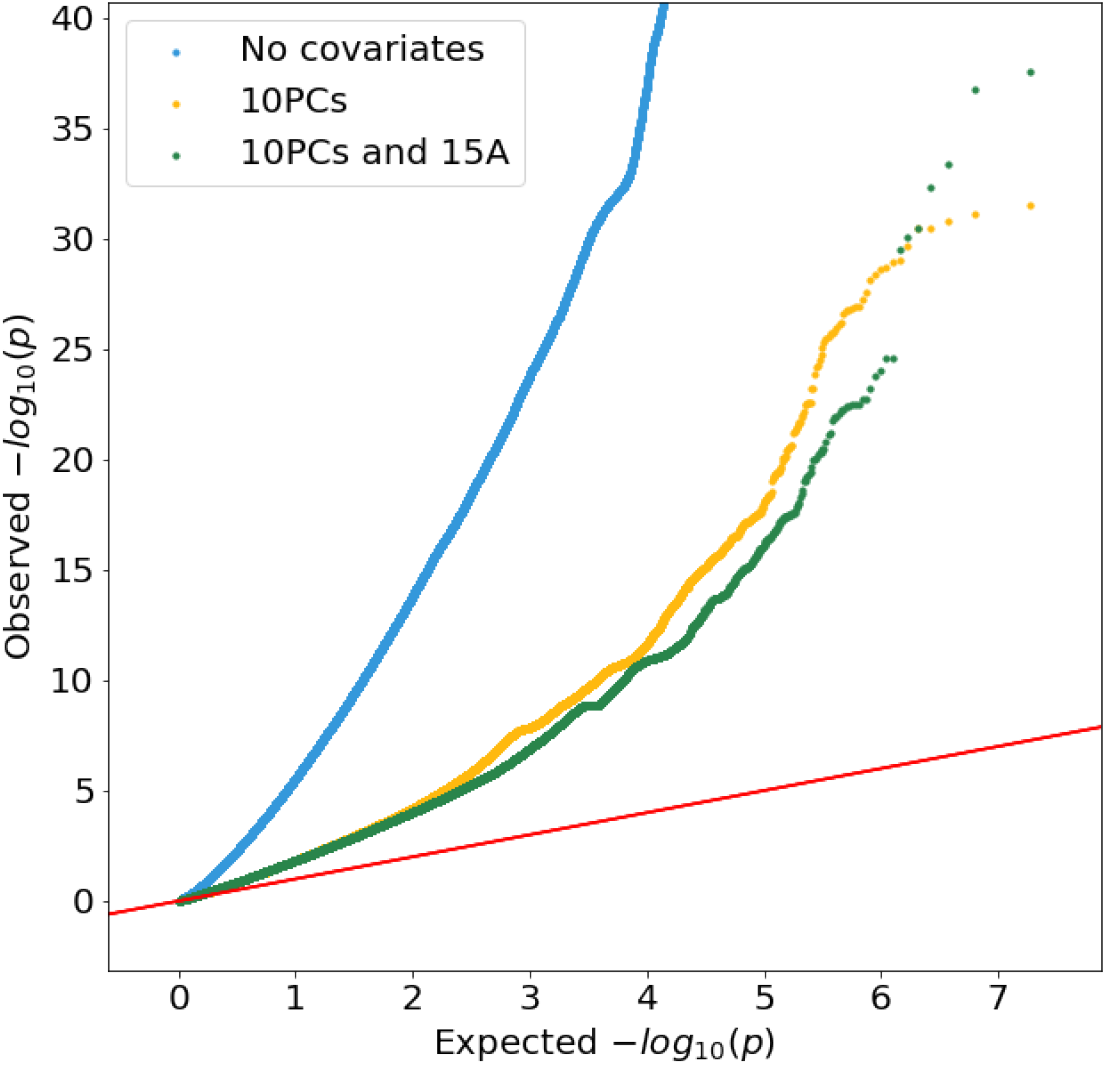
Quantile-Quantile plot (Q-Q) Q-Q plots for no covariates (blue), when adding 10 PCs (yellow) and when adding 10 PCs and 15 Archetype coefficients (green).

### Relationship between AA, K-Means, and K-Medioids

We include additional plots comparing the *K* = 3 cluster centers of AA, ADMIXTURE, K-Means, and K-Medioids in Figure 11 and *K* = 5 in Figure 12.

**Fig 11.**
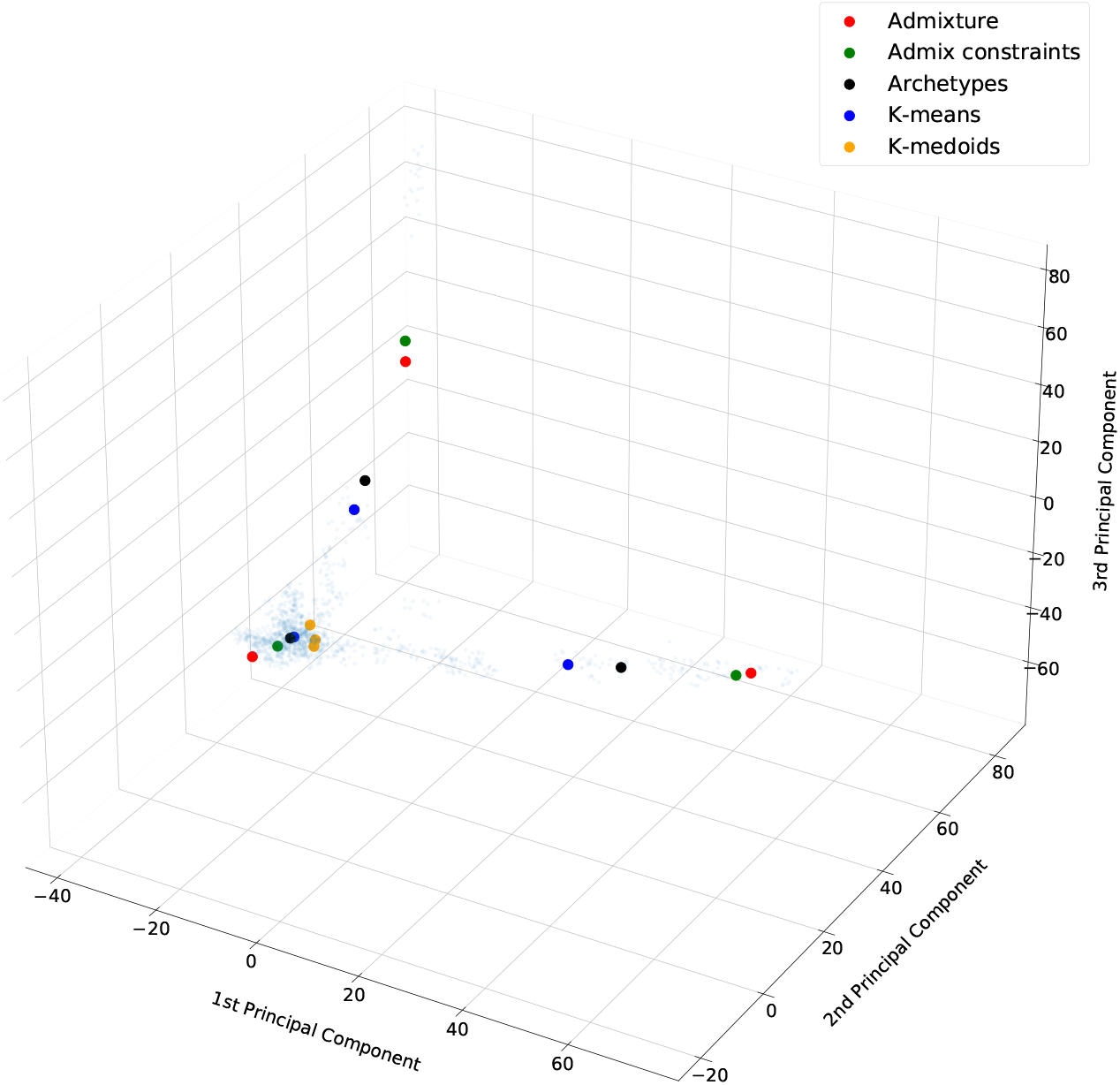
Comparison of cluster centroids from different methods. Cluster centroids learned by ADMIXTURE, ADMIXTURE with sparsity regularization, Archetypal Analysis, K-Means, and K-Medoids

**Fig 12.**
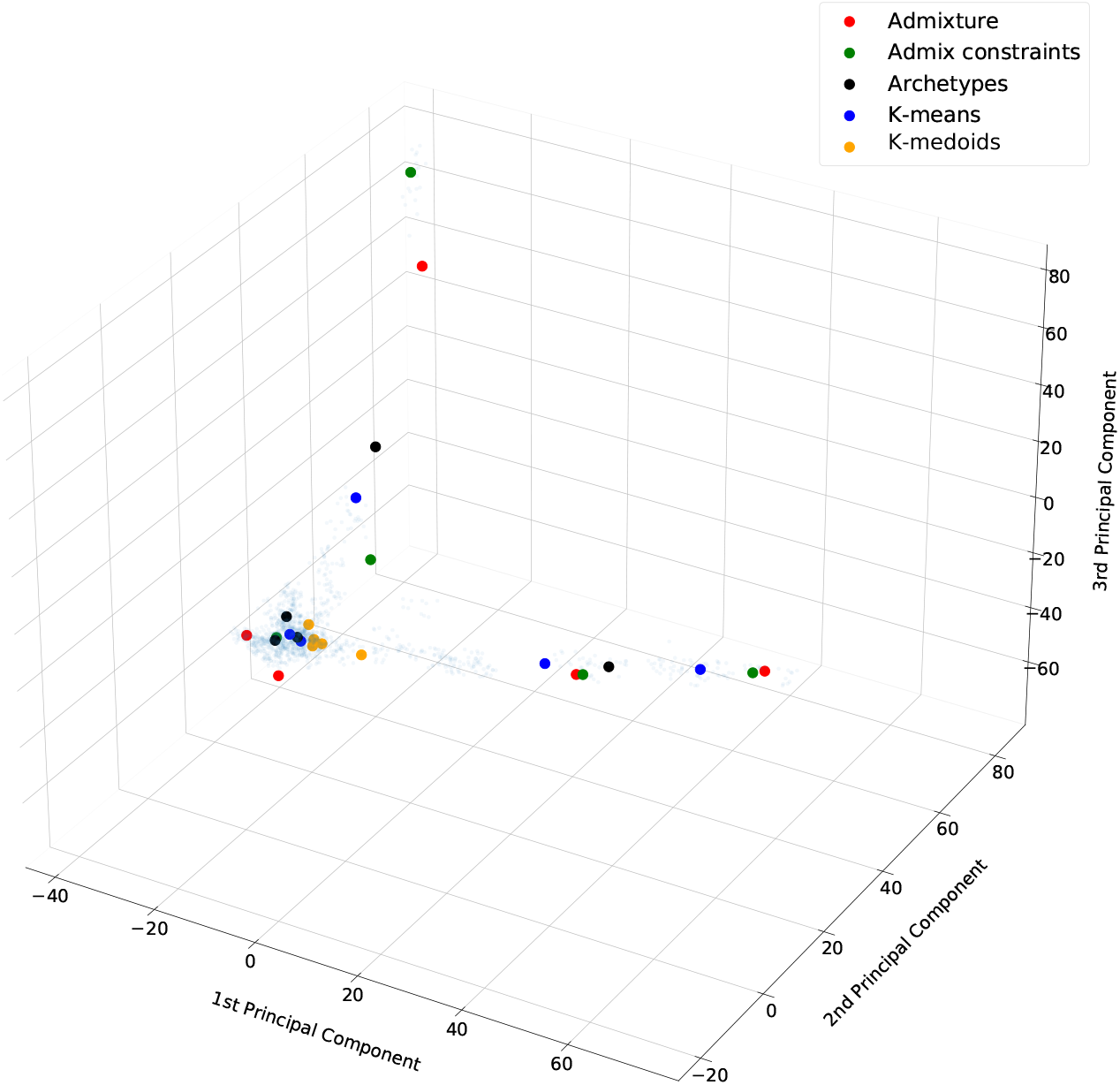
Comparison of cluster centroids from different methods. Cluster centroids learned by ADMIXTURE, ADMIXTURE with sparsity regularization, Archetypal Analysis, K-Means, and K-Medoids

